# Divergent DNA methylation signatures of X chromosome regulation in marsupials and eutherians

**DOI:** 10.1101/2020.08.26.269068

**Authors:** Devika Singh, Dan Sun, Andrew G. King, David E. Alquezar-Planas, Rebecca N. Johnson, David Alvarez-Ponce, Soojin V. Yi

**Affiliations:** School of Biological Sciences, Georgia Institute of Technology, Atlanta, Georgia, USA; Australian Museum Research Institute, Australian Museum, Sydney, New South Wales, Australia; National Museum of Natural History, Smithsonian Institution, Washington, DC, USA; Department of Biology, University of Nevada Reno, Reno, Nevada, USA

**Keywords:** X chromosome inactivation, marsupial, DNA methylation, RSX, koala, whole genome bisulfite sequencing (WGBS)

## Abstract

X chromosome inactivation (XCI) mediated by differential DNA methylation between sexes is well characterized in eutherian mammals. Although XCI is shared between eutherians and marsupials, the role of DNA methylation in marsupial XCI remains contested. Here we examine genome-wide signatures of DNA methylation from methylation maps across fives tissues from a male and female koala (*Phascolarctos cinereus*) and present the first whole genome, multi-tissue marsupial “methylome atlas.” Using these novel data, we elucidate divergent versus common features of marsupial and eutherian DNA methylation. First, tissue-specific differential DNA methylation in marsupials primarily occurs in gene bodies. Second, females show significant global reduction (hypomethylation) of X chromosome DNA methylation compared to males. We show that this pattern is also observed in eutherians. Third, on average, promoter DNA methylation shows little difference between male and female koala X chromosomes, a pattern distinct from that of eutherians. Fourth, the sex-specific DNA methylation landscape upstream of *Rsx*, the primary *lnc*RNA associated with marsupial XCI, is consistent with the epigenetic regulation of female-(and presumably inactive X chromosome-) specific expression. Finally, we utilize the prominent female X chromosome hypomethylation and classify 98 previously unplaced scaffolds as X-linked, contributing an additional 14.6 Mb (21.5 %) to genomic data annotated as the koala X chromosome. Our work demonstrates evolutionarily divergent pathways leading to functionally conserved patterns of XCI in two deep branches of mammals.

## Introduction

X chromosome inactivation (XCI) is an iconic example of sex chromosome regulation in which one of the two X chromosomes in females is silenced, a mechanism thought to adjust the expression levels of X-linked genes (1). Although XCI occurs in both eutherian and marsupial mammals (2), there are several notable differences in XCI between the two lineages. First, in eutherians, the transcription of *Xist* RNA from the inactive X chromosome is essential for XCI (3–5). However, the *Xist* locus is not present in marsupials (6, 7). Instead, another *lnc*RNA gene, *Rsx*, drives marsupial XCI (8). Second, marsupials exhibit ‘imprinted’ XCI by selectively silencing the paternal X chromosome (9, 10). In contrast, XCI in eutherians occurs randomly between the maternally and paternally derived X chromosomes, although paternal XCI has been observed during early rodent development (11, 12). Third, while eutherian XCI involves the exclusion of active histone marks and the recruitment of repressive histone marks on the inactive X chromosome (13), marsupial X chromosomes do not show a consistent pattern (10, 14). These differences suggest that evolutionary pathways leading to XCI likely differ between eutherians and marsupials, and that novel insights into the mechanism of XCI can be gained from comparative studies.

The role of DNA methylation on marsupial XCI has been particularly controversial. Immunofluorescent labeling studies observed relative hypomethylation of the inactivate X chromosome in marsupials (15, 16). Other studies found little difference in DNA methylation between active and inactive marsupial X chromosomes (10, 17, 18). Recently, Waters et al. (19) analyzed reduced representation bisulfite sequencing (RRBS) data of a male and female opossum (*Monodelphis domestica*) and proposed that female X chromosomes in marsupials, but not in eutherians, exhibit hypomethylation near the transcription start sites (TSSs). Notably, these previous DNA methylation studies either examined a small number of CpGs or employed methodologies that over-represent promoters and CpG islands (in case of RRBS, (20)). Since patterns of DNA methylation vary greatly among distinctive genomic regions with different functional consequences, it is necessary to extend our knowledge to unbiased, whole-genome assays of DNA methylation.

We used the whole-genome bisulfite sequencing, which captures the DNA methylation state of every nucleotide in the genome, of the modern koala (*Phascolarctos cinereus*) to understand the role of DNA methylation on XCI in marsupials and to resolve previous controversies. We compared DNA methylation maps of five diverse tissues from a male and a female koala, providing the first multi-tissue, whole genome methylome resource of any marsupial with information on tissue-specific variation of DNA methylation. Utilizing these novel resources, we show distinctive impacts of DNA methylation on tissue-specific gene expression in marsupials, as well as on XCI in eutherians and marsupials. We further classify previously undetected X-linked regions from this key marsupial species using characteristic features of X chromosome DNA methylation. Our findings provide new insights into the evolutionary pathways leading to functionally convergent yet mechanistically divergent pathways of XCI and regulation of gene expression in two deep branches of mammals.

## Results

### Genome-wide patterns of DNA methylation in the modern koala

We generated whole genome bisulfite sequencing (WGBS) data from five tissues (brain, lung, kidney, skeletal muscle, and pancreas) from a male (“Ben,” Australian Museum registration M.45022) and female koala (“Pacific Chocolate,” Australian Museum registration M.47723). The mean depth of coverage fell between 9.9× and 14.6× (Supplementary Table 1). Autosome-linked scaffolds displayed GC content ranging between 38.58 and 39.59%. Scaffolds linked to chromosome X had higher GC contents compared to autosomal scaffolds (Supplementary Table 2), similar to (21). We performed a hierarchical clustering analysis and observed a clear clustering of samples by tissue (Fig. 1A) with the pancreas samples exhibiting the most unique methylation signature while the kidney and lung samples share the most similar methylation profiles.

**Figure 1.**
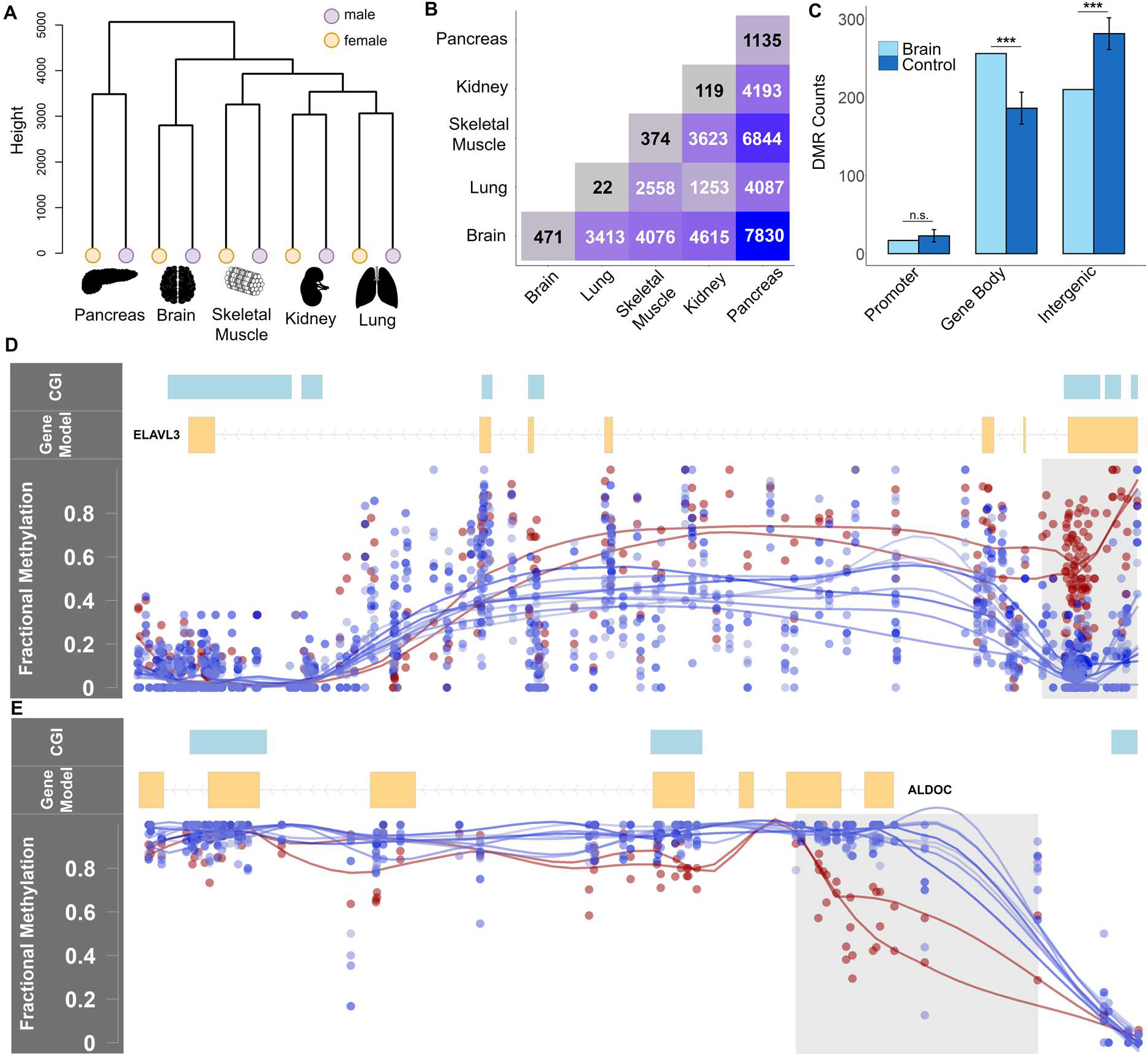
Overview of DNA methylation patterns across the koala genome. (A) Hierarchical clustering of DNA methylation of five tissues. (B) Tissue-specific and shared differentially methylation regions (DMRs) between tissues. (C) Enrichments of DMRs in each functional region in comparison to length and GC matched control regions (***p < 0.0001, n.s. Not significant, from 10,000 bootstraps) from brain samples. Error bars depict standard deviation. (D-E) Examples of two brain-specific DMRs (highlighted in grey) with corresponding CpG fractional methylation for the two brain (red) and eight remaining tissues (blue). Line smoothing performed using local regression (LOESS). (D) *ELAVL3* is a neural specific RNA-binding protein linked to the maintenance of Purkinje neuron axons (49). A 1.84 kb region containing three CpG Islands (CGI) and overlapping the first exon of *ELAVL3* was hypermethylated in the brain compared to all other tissues, and this gene was up-regulated in the brain compared to the kidneys (probability of differential expression > 98% from NOISeq). (D) *ALDOC* encodes a catalytic enzyme responsible for the conversion of fructose-1,6-bisphosphate to glyceraldehyde 3-phosphate and dihydroxyacetone phosphate. A 945 bp, brain-specific hypomethylated DMR overlaps the promoter and part of *ALDOC*’s gene body. This gene was up-regulated in brain samples compared to kidney samples (probability of differential expression > 96% from NOISeq).

### Differential DNA methylation between tissues

We identified shared and tissue-specific differentially methylated regions (DMRs) using BSmooth (22). Tissue-specific DMRs were defined as regions that were differentially methylated in a particular tissue compared to all other tissues in a pairwise analysis, while shared DMRs were those found in multiple tissues (Fig. 1B). Consistent with the results of the clustering analysis, the pancreas samples contained the most tissue-specific DMRs followed by the skeletal muscle, brain, kidney and lung (Fig. 1B). The majority (50-53%) of tissue-specific DMRs fell in gene bodies (Fig. 1C, Supplementary Fig. 1, and Supplementary Table 3), a significant excess compared to length and GC matched control regions (fold enrichment (FE) = 1.25~1.44, p < 0.0001 based on 10,000 bootstraps; Fig. 1C, Supplementary Fig. 1 and Supplementary Table 3). On the other hand, DMRs were significantly depleted in intergenic regions compared to the control regions (p < 0.05 based on 10,000 bootstraps; Fig. 1C, Supplementary Fig. 1, and Supplementary Table 3). Genes containing tissue-specific DMRs were enriched in specific biological functions (Table 1). For example, brain specific DMRs were linked to genes associated with neural developmental processes such as neurogenesis and central nervous system development.

**Table 1.**
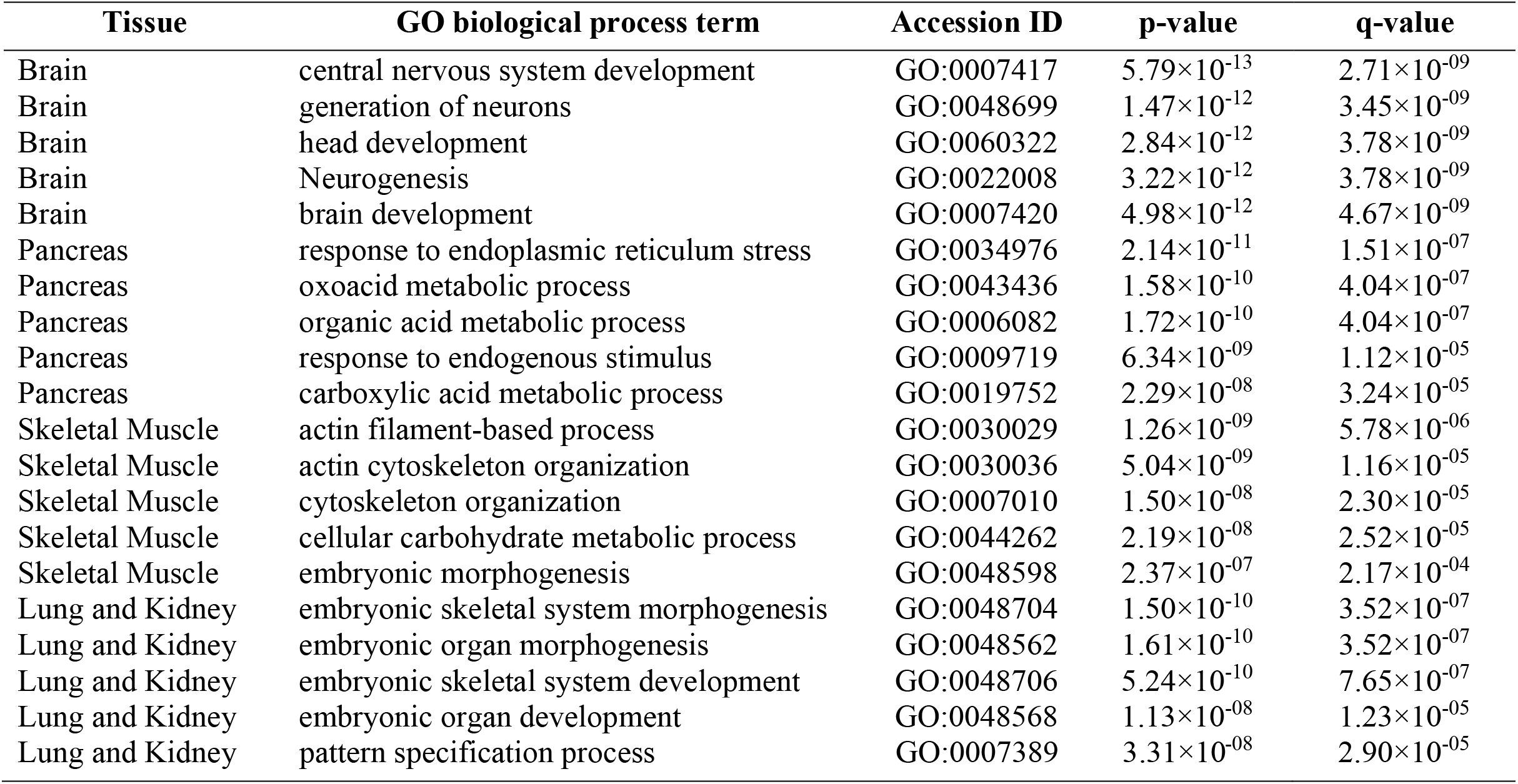
Functional annotation of enriched biological processes associated with gene sets containing tissue-specific differentially methylated regions (DMRs). Gene ontology (GO) terms are presented for the top five most significantly enriched results of each tissue after correcting for multiple testing (FDR < 0.05). As the numbers of tissue-specific DMRs for lung (n=22) and kidney (n=119) samples were so few, the corresponding gene sets were combined for this analysis.

We integrated our methylome data with a previously generated RNA-seq koala transcriptomes (23). Of the 12 tissues surveyed in that study, three tissues (kidney, brain, and lung) overlapped with the tissues examined here. Promoter DNA methylation and gene expression were significantly negatively correlated across the entire genome (Table 2, Supplementary Fig. 2). In contrast, the relationship between gene body DNA methylation and expression was complex; both extremely hypomethylated and hypermethylated gene bodies showed high gene expression (Table 2, Supplementary Fig. 2). We considered differentially methylated genes (DMGs) containing DMRs between the brain and kidney samples (n = 1944 genes from n = 4,615 DMRs) compared with differentially expressed genes (DEGs) between brain and kidney RNA-seq samples. Currently, available RNA-seq data from koalas do not include sufficient biological replicates. We overcame this limitation by simulating replicates within each RNA-seq data set (NOISeq, (24)) and identified 600 putative DEGs (probability of differential expression > 95% according to the NOISeq). We found that DMGs were significantly more likely to be differentially expressed than genes that did not contain DMRs, exhibiting a 1.54-fold enrichment (χ^2^ = 33.07, p < 0.0001). In addition, differential expression between tissues was significantly correlated with differential promoter DNA methylation between tissues (Supplementary Fig. 3A). Both relative hypomethylation and relative hypermethylation of gene bodies were associated with increased expression (Supplementary Fig. 3B). In Fig. 1D and 1E, we show DNA methylation patterns of two representative genes containing brain specific DMRs.

**Table 2.**
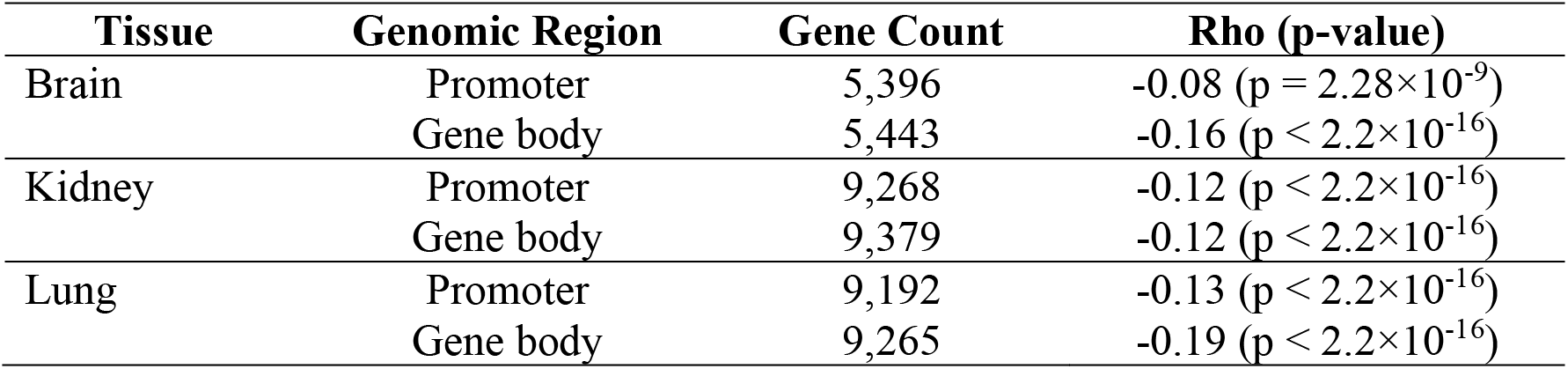
Correlation analysis of mean promoter and gene body DNA methylation and ranked gene expression. Spearman’s rank correlation coefficients (ρ) and associated significances are reported for all tissues with both whole genome bisulfite sequencing (WGBS) data and RNA-seq expression data.

| Tissue | Genomic Region | Gene Count | Rho (p-value) |
| --- | --- | --- | --- |
| Brain | Promoter | 5,396 | -0.08 ($p = 2.28 \times 10^{-9}$) |
| | Gene body | 5,443 | -0.16 ($p < 2.2 \times 10^{-16}$) |
| Kidney | Promoter | 9,268 | -0.12 ($p < 2.2 \times 10^{-16}$) |
| | Gene body | 9,379 | -0.12 ($p < 2.2 \times 10^{-16}$) |
| Lung | Promoter | 9,192 | -0.13 ($p < 2.2 \times 10^{-16}$) |
| | Gene body | 9,265 | -0.19 ($p < 2.2 \times 10^{-16}$) |

### Global hypomethylation of female X chromosome in koalas

The koala genome project used cross-species chromosome painting data to identify 24 putative X chromosome scaffolds and 406 putative autosomal scaffolds (25). As expected from 2:1 ratio of X chromosomes in females compared to males, the median depth of coverage of CpGs on the putative X scaffolds were consistently higher (approximately 2-fold) in female samples compared to male samples (p < 2.2 × 10^−16^, Mann-Whitney *U* test, Supplementary Fig. 4A). Additionally, the proportion of reads mapped to the putative X scaffolds showed a distinct bimodal distribution whereby the male samples cluster close to 1.3% and the female samples cluster near 2.4% (Supplementary Fig. 4B). In contrast, male and female samples were indistinguishable with respect to read mapping to putative autosomes (Supplementary Fig. 4D).

The global DNA methylation level of the female X chromosome was lower than that of the male X chromosome in all koala tissues examined (Fig. 2A, B and Supplementary Fig. 5, p < 2.2 × 10^−16^, Mann-Whitney *U* test). A comparison to autosomal DNA methylation indicated that the X chromosome exhibits reduced DNA methylation in females. Consequently, we use the term ‘female hypomethylation’ (as opposed to male hypermethylation) (Fig. 2C) consistently in this work. As a comparative basis of the DNA methylation patterns observed in eutherian mammals, we performed parallel analyses on human DNA methylation data (methods). Human X chromosomes were also globally hypomethylated in females compared to males (Fig. 2A, C). Female hypomethylation was observed in all functional regions across the X chromosome (Fig. 2D), but most pronounced in gene bodies and intergenic regions. Promoters showed the least difference between males and females. As expected, the autosomal scaffolds did not display a significant variation between female and male methylation levels in any functional region (Fig. 2E). Fig. 2F depicts the DNA methylation difference between male and female X chromosomes in humans. In contrast to the pattern observed in koalas, promoters on the human female X chromosomes were hypermethylated compared to those on the male X chromosome, congruent with previous studies (26–28).

**Figure 2.**
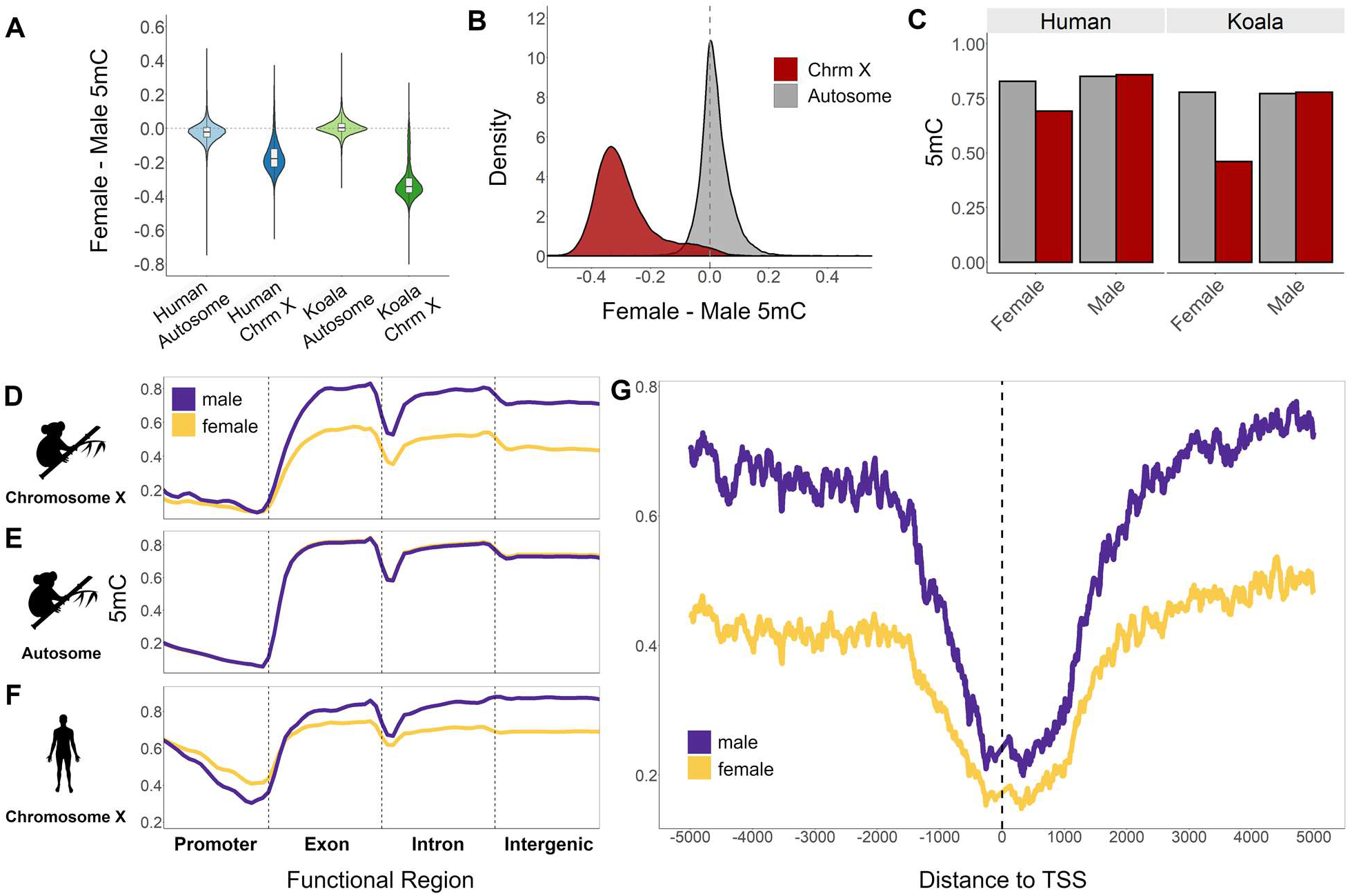
Global patterns of female and male DNA methylation (5mC) in eutherian and marsupial X chromosomes. (A) Comparison of chromosome-wide DNA methylation between sexes in human and koala brains. (B) Distributions of the DNA methylation difference between female and male koalas in autosomes and the X chromosome. (C) Comparing fractional DNA methylation between males and females demonstrate that the female X chromosomes are hypomethylated in both humans and koalas. (D-F) DNA methylation differences between males and females of (D) the X chromosome and (E) autosomes of koalas, and (F) the X chromosome of humans for all male (purple) and female (orange) samples. Line smoothing was performed using local regression (LOESS). (G) Average fractional methylation of CpGs in 100-bp sliding windows using a 10 bp step size in a 5 Kb region upstream and downstream of all chromosome X linked gene’s transcription start sites (TSSs) across koala tissues.

### Promoter DNA methylation is not a universal driver of sex-specific expression in koalas

To investigate the implications of the observed sex-specific DNA methylation, we once again utilized the published RNA-seq koala transcriptomes (23). Of the total RNA-seq dataset, only one tissue (kidney) had expression data from both a male and female koala that could be directly compared to our methylome dataset. Consequently, the other tissues were not considered for further analysis. We calculated the log-transformed fold change of female and male gene expression values using NOISeq, capitalizing on its ability to simulate technical replicates. Of the 209 X-linked genes, 36 (17.2%) exhibit female overexpression while 11 (5.3%) show male overexpression (probability of differential expression > 95% based on NOISeq, Supplementary Fig. 6A). Although, on average, autosomal genes also exhibited slight female-bias of expression (Supplementary Fig. 6B, C), the increase was more substantial in the X chromosome (mean chromosome X female to male log2 fold change = 0.50, autosome female to male expression log2 fold change = 0.24). We calculated the female and male fractional methylation difference in X chromosome-linked promoters and gene bodies correlated with gene expression difference (N = 209 gene bodies and N = 206 promoters, excluding 3 promoters with CpGs coverage < 3). In promoters, no significant relationship was observed (Supplementary Fig. 7A). In fact, the proportions of significantly female-over expressed genes were similar between female-hypermethylated promoters and female-hypomethylated promoters (Supplementary Fig. 7B). Interestingly, female and male DNA methylation difference in gene bodies showed an overall negative correlation with gene expression (Spearman’s rank correlation coefficient, ρ = −0.14, p = 0.04, Supplementary Fig. 7C). A deeper analysis of the relative DNA methylation and expression levels revealed that both extreme hypomethylation and hypermethylation are associated with increased expression (Supplementary Fig. 7C) attesting to the complexity and heterogeneity of the relationship between gene body DNA methylation and gene expression.

### The *Rsx* region displays a pattern suggesting methylation driven control of X chromosome regulation in koalas

Sex-specific DNA methylation of a key regulator of XCI in other marsupials, the *lnc*RNA gene *Rsx*, has been associated with its differential expression (8, 10). Based on the sequence homology with the *Rsx* gene from the gray short-tailed opossum (*Monodelphis domestica*) (8), we identified a 29.8 Kb candidate *Rsx* sequence (Methods), using PacBio long read sequencing by Johnson et al. (25). Interestingly, we observed a female hypomethylated region containing two CpG islands upstream of the candidate *Rsx* covering 101 CpGs exhibiting a 36% reduction in mean female DNA methylation compared to mean male DNA methylation (mean sex difference: −0.36 ± 0.14, Fig. 3). The mean DNA methylation difference between sexes within this region was in the top 23% of the distribution of differences across all X-linked scaffolds of the combined tissues. The expression of the *lnc*RNA annotated within the candidate *Rsx* gene region was significantly greater in females (mean normalized read count = 6987.1) than in males (mean normalized read count = 16, p < 0.05 from DeSeq2 using the Wald test). When considering a subset of tissue expression data with matched male and female samples (spleen, kidney, and lung), the results remained robust across different tools to measure differential gene expression (Table 3).

**Figure 3.**
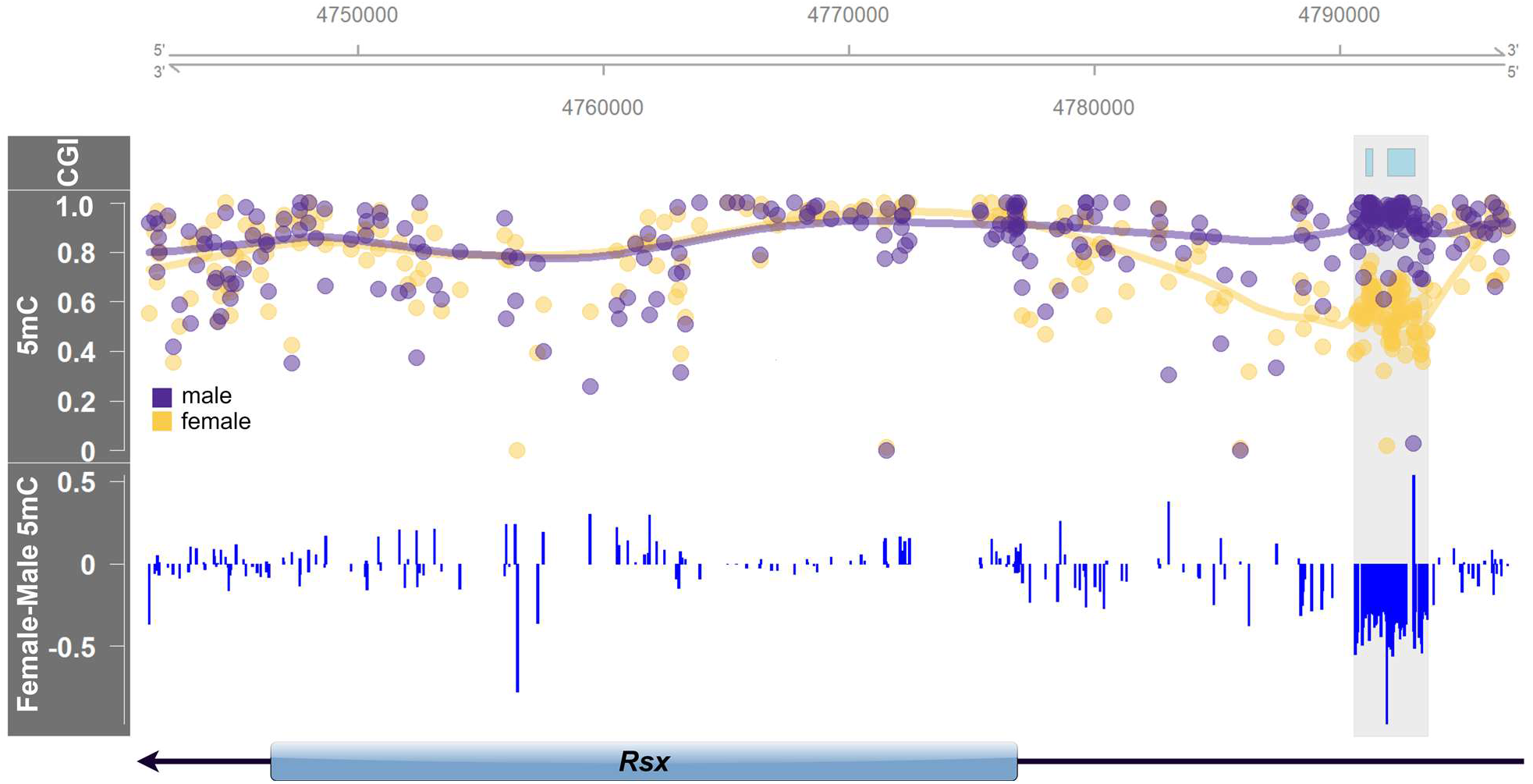
Annotation of genomic of DNA methylation (5mC) around *Rsx*. The top panel identifies CpG islands (CGI), the middle panel reports the absolute male (purple) and female (orange) fractional methylation at each CpG, and the bottom panel shows the female and male fractional methylation difference. Highlighted in grey across all panels is the female hypomethylated region upstream of the *Rsx* TSS.

**Table 3.** Sex-based differential expression of the *lnc*RNA *Rsx* utilizing different data subsets and expression quantification tools. Normalized expression count values and significance of sex-based differential expression is shown for three data subsets using two expression quantification tools. All data refers to the dataset considering all 15 RNA-seq samples (7 male and 8 female). Matched data includes the tissues with both male and female RNA-seq samples (brain, kidney, and lung), and the kidney data is reported independently. DeSeq2 reports significance as an associated p-value from the Wald test while NOISeq reports a probability of differential expression threshold.

| Expression Dataset | Tool | Female Count | Male Count | Significance |
| --- | --- | --- | --- | --- |
| All Data (n = 15) | DeSeq2 | 6987.1 | 16 | p-value = 0.05 |
| Matched Data (n = 6) | DeSeq2 | 6837.6 | 0 | p-value = $2.04 \times 10^{-30}$ |
| Matched Data (n = 6) | NOISeq | 7872.4 | 0.67 | Probability = 99.99% |
| Kidney Data (n = 2) | NOISeq | 4074.4 | 0.68 | Probability = 99.99% |

### Identification of novel candidate X-linked scaffolds by sex-specific methylation patterns

We have demonstrated above that X-linked scaffolds exhibit several distinctive WGBS patterns in koalas, including different depths of coverage per CpG, distinctive clustering based on the proportion of mapped reads, and unique methylation distributions between females and males (Supplementary Fig. 4). We thus quantified DNA methylation differences between females and males to determine if additional candidate X scaffolds existed within the 6.7% of the koala assembly that remained unclassified. We identified an additional 98 scaffolds that showed a clear shift towards female hypomethylation (mean female-male 5mC for all candidate X scaffolds was −0.25 ± 0.12) (Supplementary Tables 4,5). These candidate scaffolds contributed 14.6 Mb (21.5%) to the total genomic region annotated as the koala X chromosome. All candidate scaffolds followed the expected bimodal distribution seen in the putative X scaffolds; male samples clustered around 0.3% and the female samples clustered around 0.5% (Supplementary Fig. 4C). This clustering pattern could be attributed to the 2:1 ratio of X chromosomes in females compared to males that was not observed in putative autosomes.

## Discussion

In this study, we analyzed novel nucleotide-resolution genomic DNA methylation maps of five tissues of a male and a female koala. The overall DNA methylation levels of koala tissues are on par with those in other mammals (27, 29, 30), and show heavy genome-wide DNA methylation punctuated by hypomethylation of CpG islands (e.g., Fig. 1D,E). Tissue-specific differential DNA methylated regions (DMRs) were significantly enriched in gene bodies. Gene body methylation is an ancestral form of DNA methylation in animal genomes (e.g., (31, 32)), yet its role in regulation of gene expression is less well understood than promoter DNA methylation owing in part to the heterogeneity of gene body regions with respect to sequence length, composition and functionality of introns and exons, and CpG density compared to promoter regions (31, 33). For example, DNA methylation levels of the first exons/introns of genes are negatively correlated with gene expression (34–36), and tend to be different from downstream genic regions (35). On the other hand, high levels of cumulative gene body DNA methylation is positively correlated with gene expression and may reduce spurious transcription of intragenic RNA (37, 38). These studies illustrate the potential for both positive and negative regulatory control of gene expression by gene body DNA methylation.

We show that female X chromosomes are globally hypomethylated compared to both the male X chromosome and the autosomes of both sexes (Fig. 2). Even though it may appear counterintuitive at the first glance, the hypomethylation of female X chromosome appears as a common feature of eutherian and marsupial mammals. Hellmann and Chess (26) showed that the inactive X chromosomes of humans had reduced gene body DNA methylation. Recent whole genome bisulfite sequencing analyses of mouse (27) and humans (28) showed that the hypomethylation of inactive X chromosome is pervasive in gene bodies and intergenic regions (Fig. 4). Waters et al. (19) performed reduced representation bisulfite sequencing (RRBS) of several species including mouse and opossum, and concluded that the reduction of methylation in gene bodies was specific to marsupials, but not in eutherians (19). The reason why they did not observe methylation difference in mouse might be due to the inherent bias of RRBS, which disproportionately samples regions with high GC content (20). High GC-content regions tend to be hypomethylated (39, 40) and show less variation of DNA methylation. For example, the degree of methylation difference between sexes in koala promoters is small compared to other genomic regions, at least partly due to high GC content of promoters (Supplementary Fig. 8). How chromosome-wide DNA hypomethylation is linked to chromosome-wide gene silencing is currently unknown. Interestingly, marsupial genomes harbor an additional copy of *DNMT1* (41), which could lead to functional divergence between the mammalian lineages. Analyses of DMNT expression however did not indicate significant differential expression of *DNMT*s between sexes (probability of differential expression using NOISeq < 95%). Additional analysis with broad and balanced sampling of both sexes is necessary to rigorously test the impact of the marsupial *DNMT1* duplication on gene expression.

**Figure 4.**
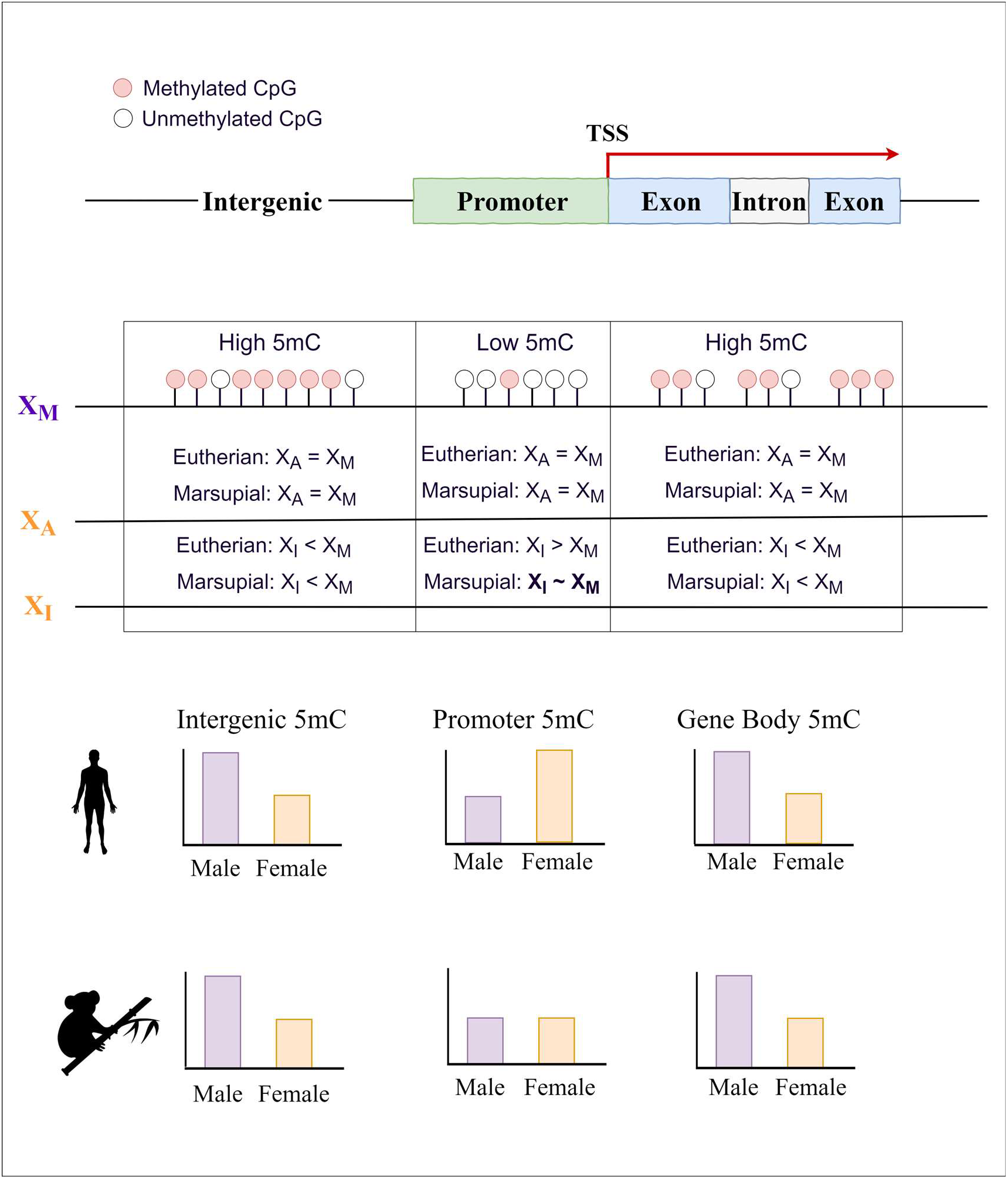
Model of DNA methylation (5mC) patterns across genomic functional regions for eutherian mammals and marsupials. Eutherian mammals exhibit increased DNA methylation of promoter regions and CpG islands coupled with a relative depletion of DNA methylation across gene bodies and intergenic regions on the inactive X chromosome (X_I_) compared to the active X chromosome (X_A_) in females. Marsupial mammals share similar DNA methylation depletion in gene bodies and intergenic regions of the inactive X chromosome; however, they diverge from eutherian mammals in their promoter methylation patterns. Marsupial promoters are modestly hypomethylated in the female X chromosomes (X_A_ and X_I_) compared to the male X chromosome (X_M_).

Interestingly, DNA methylation signatures of *Rsx,* the major player in XCI initiation in marsupials (8), suggest that koala *Rsx* expression is regulated by promoter CpG island DNA methylation (Fig. 3). Wang et al. (10) also showed differential DNA methylation of *Rsx* promoter in opossum. Therefore, regulation of the key initiator of XCI via DNA methylation is another parallel feature between eutherians and marsupials Notably, a recent study (42) found that *Xist* and *Rsx* harbor non-linear sequence similarity. Consequently, their shared functionality may be partially due to characteristics of tandem repeat regions.

In summary, we show that gene body DNA methylation is an important contributor to differential expression between tissues in koala. We also show that the global hypomethylation of female X chromosome is a conserved feature of X chromosome regulation in eutherian and marsupial mammals. However, X chromosome promoter methylation and the subsequent effect on the regulation of gene expression appear to be divergent between these two lineages. Regulation of the *Rsx*, on the other hand, is supported by DNA methylation, which mirrors the regulation of the eutherian *Xist* locus. Together, these conclusions illuminate the intricate evolutionary pathways that have diverged and converged to influence X chromosome regulation, XCI, and dosage compensation in eutherian and marsupial mammals.

## Methods

### Whole genome bisulfite sequencing and processing

All genomic DNA was extracted using a Bioline Isolate II Genomic DNA Extraction Kit (Cat#. BIO-52067) following the recommended protocol with an additional DNAse free RNaseA (100mg/ml) (Qiagen cat. #19101) treatment before column purification. 20mg tissue samples from brain, kidney, lung, skeletal muscle, and pancreas from a female koala, “Pacific Chocolate” (Australian Museum registration M.45022), and a male koala, “Ben” (Australian Museum registration M.47723), were supplied to the Ramaciotti Centre for Genomics for methylome sequencing. The bisulfite conversion was carried out by using the EX DNA Methylation-Lightning Kit (Zymo cat. #D5030) and the WGBS libraries were constructed using the TruSeq DNA methylation kit (Illumina cat.# EGMK81213). The libraries were sequenced on a NovaSeq6000 S2 (Illumina) using the 2 × 100bp PE option. Information on samples, coverage, and read counts are provided in Supplementary Table 1. Processing of the WGBS data followed previous studies (30). Bisulfite conversion rates were estimated for each WGBS sample following the methodology of methPipe's bsrate (43) (Supplementary Table 1). Strand-specific methylation calls were combined, and all samples were filtered to remove CpGs covered by fewer than three reads.

### Analyses of tissue differentially methylated regions

A hierarchical clustering tree was drawn using the fractional methylation profiles of all ten samples using hclust from R’s stats package. The distance matrix was calculated using Euclidean distances and the agglomeration method used was Ward’s method. The data for the final tree was visualized using R’s dendextend package (44). Bismark generated CpG reports were filtered to remove scaffolds that were less than 2 Mb in length, retaining 3.03 × 10^9^ (94.8%) of the genome. DMRs were called using BSmooth (22), with additional filtering of a minimum fractional methylation difference of 0.3 (30%) and at least 5 CpG sites. DMRs were considered shared between tissues if they overlapped by at least 50%. Using koala gene annotations from Ensembl (Phascolarctos_cinereus.phaCin_unsw_v4.1.97 release), promoters were defined as regions located 1000 bp upstream of the identified transcription start site (TSS). To test the enrichment of tissue-specific DMRs in functional regions, we generated 10,000 length and GC content matched controls for all unique DMRs. The lung and kidney samples shared the most similar methylation profiles and consequently had few tissue-specific DMRs (Fig. 1C). Because of this feature, the corresponding gene sets were combined for these two tissues. Functional annotation and GO term enrichment analysis was performed utilizing the ToppGene Suite (45).

### Differential DNA methylation between sexes

Johnson et al. (25) used cross-species chromosome painting data and linked 406 scaffolds spanning 2.9 Gb of sequence data to autosomal scaffolds from chromosomes 1-7 and 24 scaffolds covering 68 Mb of sequence data to chromosome X, leaving 6.7% of the genome unclassified. To explore the same relative amount of genomic space, we randomly sampled a subset of the autosomal scaffolds that were length matched with the X chromosome scaffolds, which we called the “matched autosome” dataset. These scaffolds were divided into 10-kb bins and the difference in mean fractional methylation at each 10-kb bin was compared between male and female samples for all tissues. For the analysis of human data, we used WGBS fractional methylation reports from a male brain (Epigenome ID: E071) and a female brain (Epigenome ID: E053) and the human known gene annotations from Ensembl (hg19 release). Due to its similarity in size to the human X chromosome, we used data from human chromosome 8 as our representative autosome in the comparative analysis.

### Identification of candidate X-linked scaffolds

To isolate candidate X-linked scaffolds from the 1,477 unclassified koala scaffolds, we binned the unclassified scaffolds into 10-kb windows and calculated the mean fractional methylation of the associated CpGs. We then determined the average female and male methylation differences across the bins and plotted the density of the differences for all five tissues. SVY and DS proceeded to independently select scaffolds that exhibited a statistically significant shift towards female hypomethylation from zero. The scaffolds that showed significant female hypomethylation in all five tissues and were selected by both SVY and DS were isolated (n = 98 covering 14.6 Mb of sequence with mean female-male 5mC = −0.25 ± 0.12). As an external validation, the percent of reads mapping to the putative X-linked and autosome-linked scaffolds over the total number of mapped reads was computed for the male and female sample in all tissues.

### Annotation of the koala *Rsx*

To annotate the genomic region around *Rsx*, we downloaded the published genome sequence fasta files for the partial assembly of *Rsx* from the gray short-tailed opossum (8) and the complete PacBio assembly of the koala *Rsx* (25, 42). We used BLASTN 2.2.29 (46) to align both sequences to the koala reference genome (phaCin_unsw_v4.1) and obtained genomic coordinates. The entire assembled koala *Rsx* sequence aligned with 100% identity and no gaps. Only one 30.4 kb transcript, a novel *lnc*RNA, overlapped with the annotated *Rsx* region (overlap > 90% of transcript) and was used to evaluate gene expression.

### Analysis of differential gene expression

All RNA-seq expression data used in this analysis were obtained from the previously published koala transcriptomes from eight females and seven males (23). To process the acquired data, we followed the protocol outlined by Pertea et al. (47), using the koala GTF annotation from Ensembl (Phascolarctos_cinereus.phaCin_unsw_v4.1.97.gtf.gz release) to assemble mapped reads into transcripts using StringTie 2.0 (47) with the -e-b--A <gene_abund.tab> flags. We used StringTie’s functionality for *de novo* transcript assembly to identify candidate *Rsx* transcripts. An updated GTF annotation was generated including novel transcripts using the --merge flag and then the previously generated mapped reads were reassembled into transcripts guided by this GTF file. DeSeq2 1.22.2 (48) was used to perform differential gene expression analysis between males and females. NOISeq 2.26.1 (24) was used for differential expression analysis due to its ability to simulate technical replicates within given RNA-seq data sets when no replicates are available.

## Data accessibility

The raw and processed methylation datasets generated in this study have been deposited and accessible through GEO Series accession number GSE149600 at https://www.ncbi.nlm.nih.gov/geo/query/acc.cgi?acc=GSE149600.

## Authors’ contributions

D.S., D.A.P. and S.V.Y. formed the research design; D.A.P., A.G.K., D.E.A.P, and R.N.J. generated the data; D.S., D.S. and S.V.Y. performed the analysis and wrote the initial draft; all authors contributed to and approved the final manuscript.

## Supplementary Tables

**Supplementary Table 1.** Overview of whole genome bisulfite sequencing (WGBS) data for all 10 koala samples.

| Sample | Australian Museum registration | Name | Tissue | Mapped Reads | De-duplicated Reads | Total CpGs | Coverage (%) | Mean Depth | % Reads Cov > 3× | Bisulfite Conversion Rate |
| --- | --- | --- | --- | --- | --- | --- | --- | --- | --- | --- |
| WGM145_01_S1 | M.45022 | Pacific Chocolate | Brain | 602055711 | 228244836 | 16761785 | 97.5 | 13.99 | 93.4 | 98.2 |
| WGM145_04_S2 | M.47723 | Ben | Brain | 677593548 | 233893124 | 16761785 | 97.6 | 14.61 | 93.8 | 98.0 |
| WGM145_06_S3 | M.45022 | Pacific Chocolate | Kidney | 559911229 | 213274030 | 16761785 | 97.2 | 13.62 | 92.5 | 98.7 |
| WGM145_08_S4 | M.47723 | Ben | Kidney | 589121434 | 202094771 | 16761785 | 97.2 | 12.93 | 92.3 | 98.7 |
| WGM145_09_S5 | M.45022 | Pacific Chocolate | Lung | 637878987 | 210002505 | 16761785 | 97.5 | 13.8 | 93.2 | 98.6 |
| WGM145_12_S6 | M.47723 | Ben | Lung | 663204935 | 198555030 | 16761785 | 97.3 | 12.54 | 92.4 | 98.6 |
| WGM145_14_S7 | M.45022 | Pacific Chocolate | Skeletal Muscle | 592423486 | 168450992 | 16761785 | 96.7 | 10.93 | 90.0 | 98.7 |
| WGM145_16_S8 | M.47723 | Ben | Skeletal Muscle | 605979530 | 168220022 | 16761785 | 96.8 | 11.12 | 90.4 | 98.6 |
| WGM145_19_S9 | M.45022 | Pacific Chocolate | Pancreas | 563288200 | 159866998 | 16761785 | 96.2 | 9.857 | 88.0 | 98.6 |
| WGM145_20_S10 | M.47723 | Ben | Pancreas | 598508860 | 166573663 | 16761785 | 96.6 | 10.56 | 89.4 | 98.7 |

**Supplementary Table 2.**
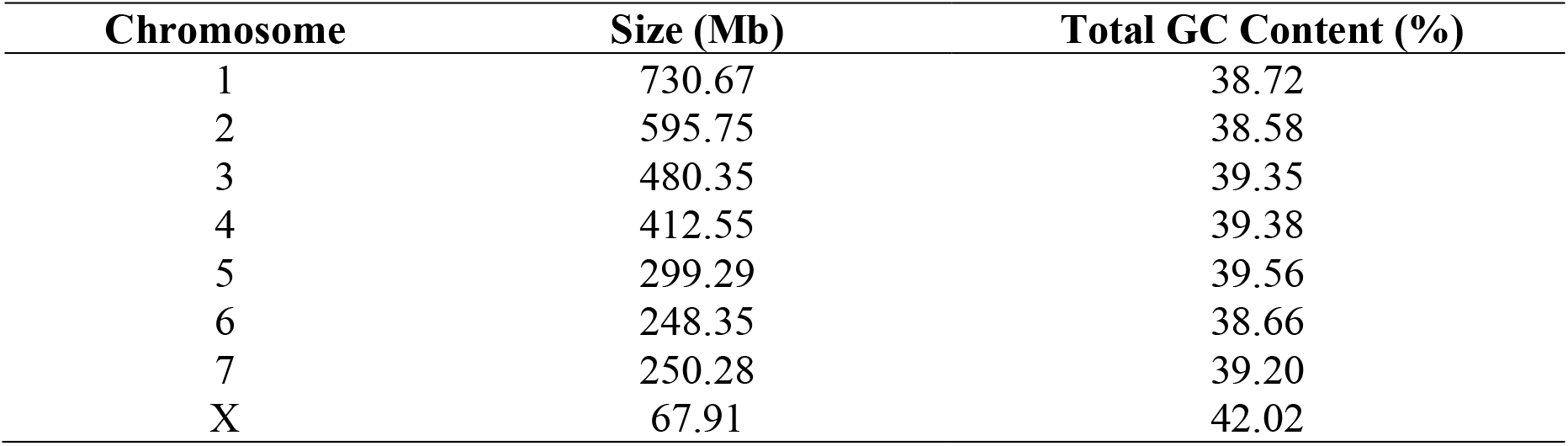
GC-content for all scaffolds linked to annotated autosomes (chromosomes 1-7) and the X chromosome.

**Supplementary Table 3.**
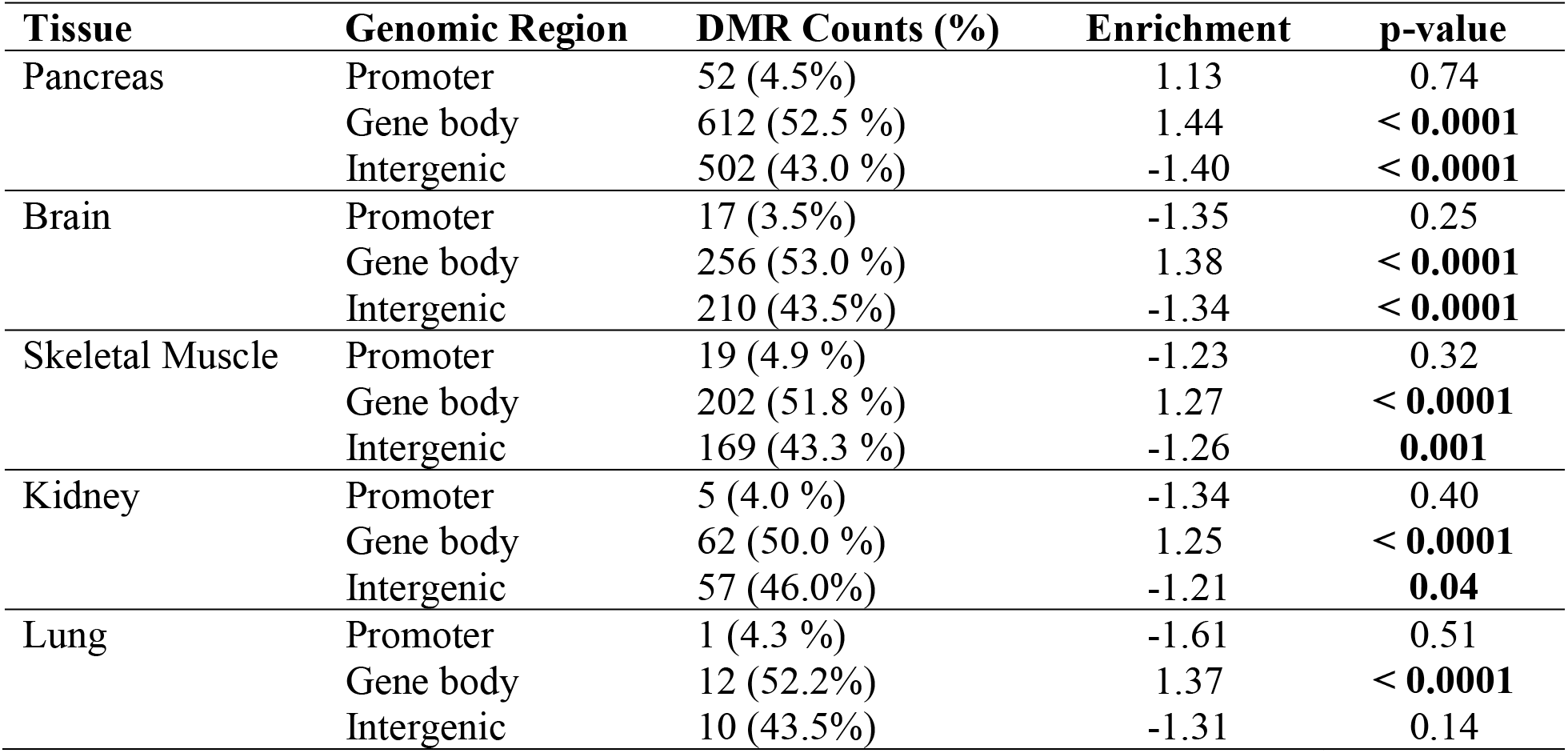
Enrichment and significance of all tissue specific DMRs compared to length and GC matched control regions. Reported are the total counts of tissue specific differentially methylated regions (DMRs) falling within one of three annotated genomic regions: promoters, gene bodies, and intergenic regions. The enrichment of DMRs in each functional region is shown through a fold change comparison with a control dataset generated from 10,000 bootstraps using length and GC matched control regions. All significant p-values (p < 0.05) are highlighted in bold.

**Supplementary Table 4.** Mean and median sex-based DNA methylation difference calculated for all candidate X-scaffolds (n=98) by tissue. The female and male mean fractional DNA methylation (methylated reads/total reads per CpG) was calculated for all CpGs within 10 kb bins across candidate scaffold.

| **Tissue** | **Median Male-Female 5mC** | **Mean Male-Female 5mC** |
| --- | --- | --- |
| Pancreas | -0.2282 ± 0.15 | -0.2066 ± 0.12 |
| Brain | -0.3273 ± 0.14 | -0.2911 ± 0.12 |
| Skeletal Muscle | -0.2748 ± 0.15 | -0.2431 ± 0.12 |
| Kidney | -0.2850 ± 0.15 | -0.2567 ± 0.12 |
| Lung | -0.2706 ± 0.14 | -0.2437 ± 0.11 |
| Combined Tissues | -0.2773 ± 0.15 | -0.2484 ± 0.12 |

## Supplementary Figures

**Supplementary Figure 1.**
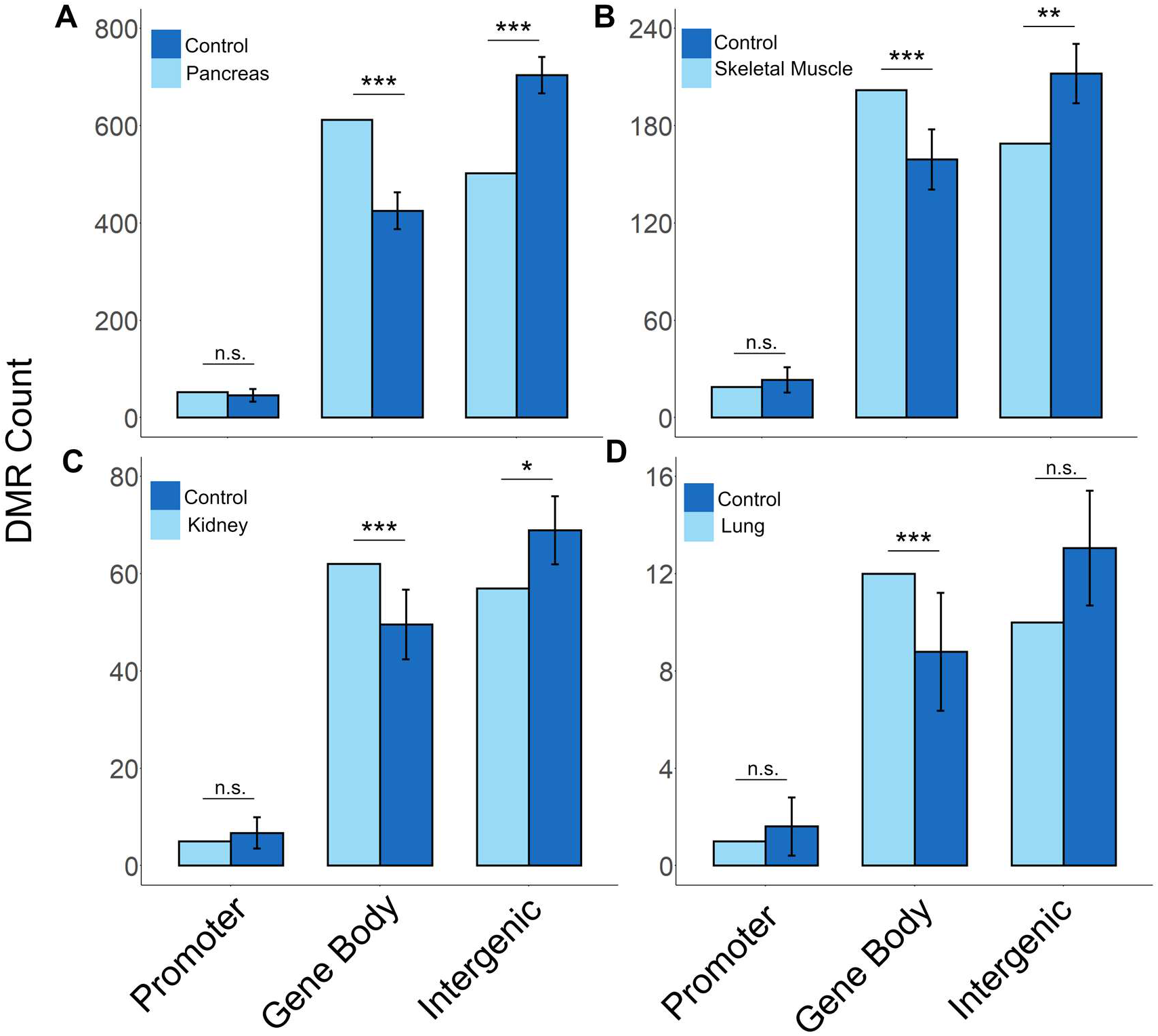
Enrichment of tissue specific differentially methylated regions (DMRs) falling within genomic functional regions. Data shown for (A) pancreas, (B) skeletal muscle, (C) kidney, and (D) lung. The enrichment of DMRs in each functional region (promoter, gene body, and intergenic regions) is shown through a comparison with length and GC matched control regions (*** indicates p < 0.0001, ** indicates p < 0.001, * indicates p < 0.05, and non-significance is shown by n.s. based on 10,000 bootstraps). Error bars indicate standard deviation.

**Supplementary Figure 2.**
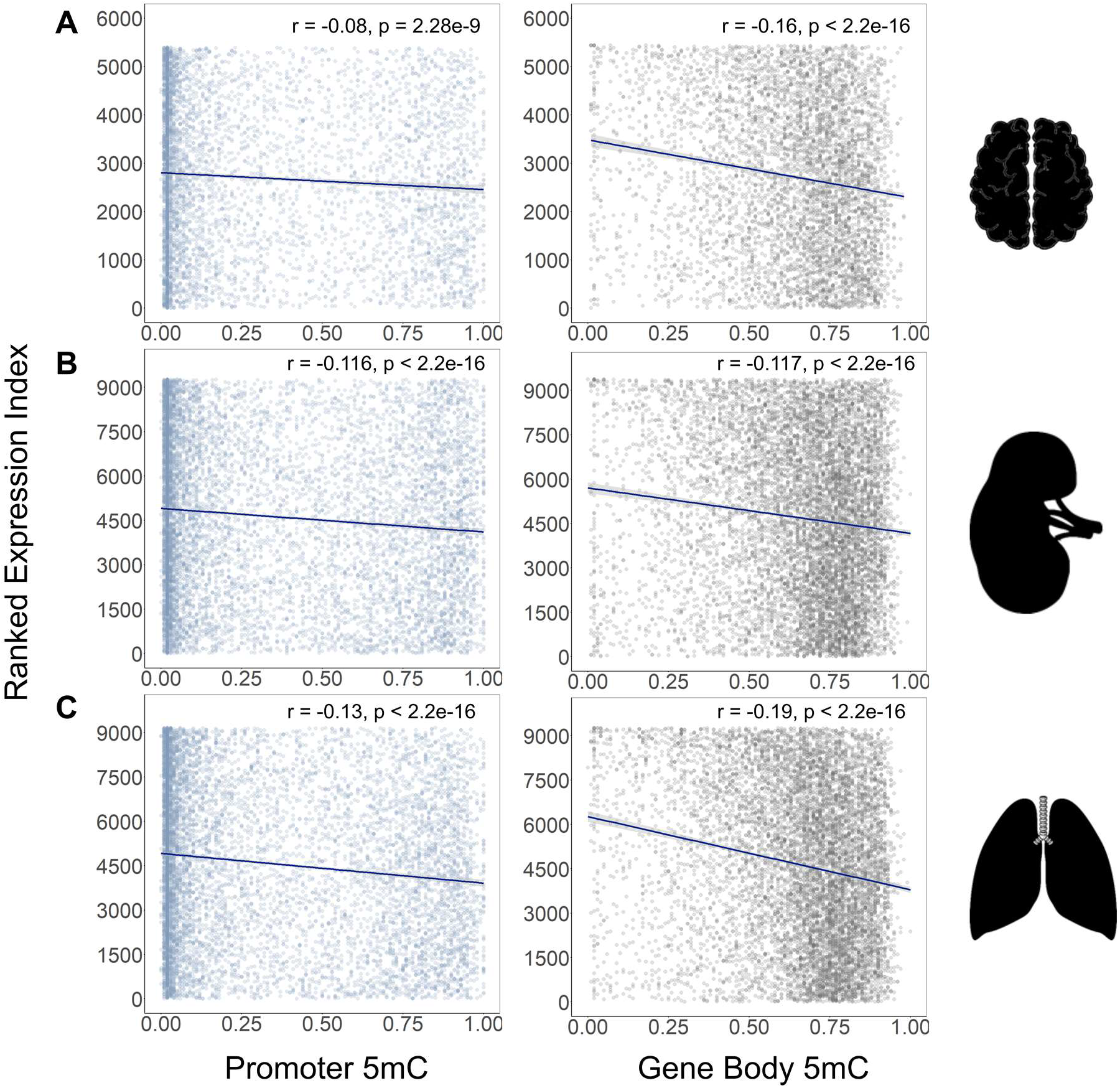
Correlation of gene expression and DNA methylation (5mC) in CpGs across promoters and gene bodies. Three tissues with both whole genome bisulfite sequencing (WGBS) DNA methylation data and RNA-seq gene expression data are shown, (A) brain (n = 5,396 promoters and n = 5,443 gene bodies), (B) kidney (n = 9,268 promoters and n = 9,379 gene bodies), and (C) lung (n = 9,192 promoters and n = 9,265 gene bodies). For A-C, TPM expression values were ranked from low to high for each gene and correlated with mean fractional DNA methylation (methylated reads/total reads per CpG site). Spearman’s rank correlation coefficients and the associated p-values are reported.

**Supplementary Figure 3.**
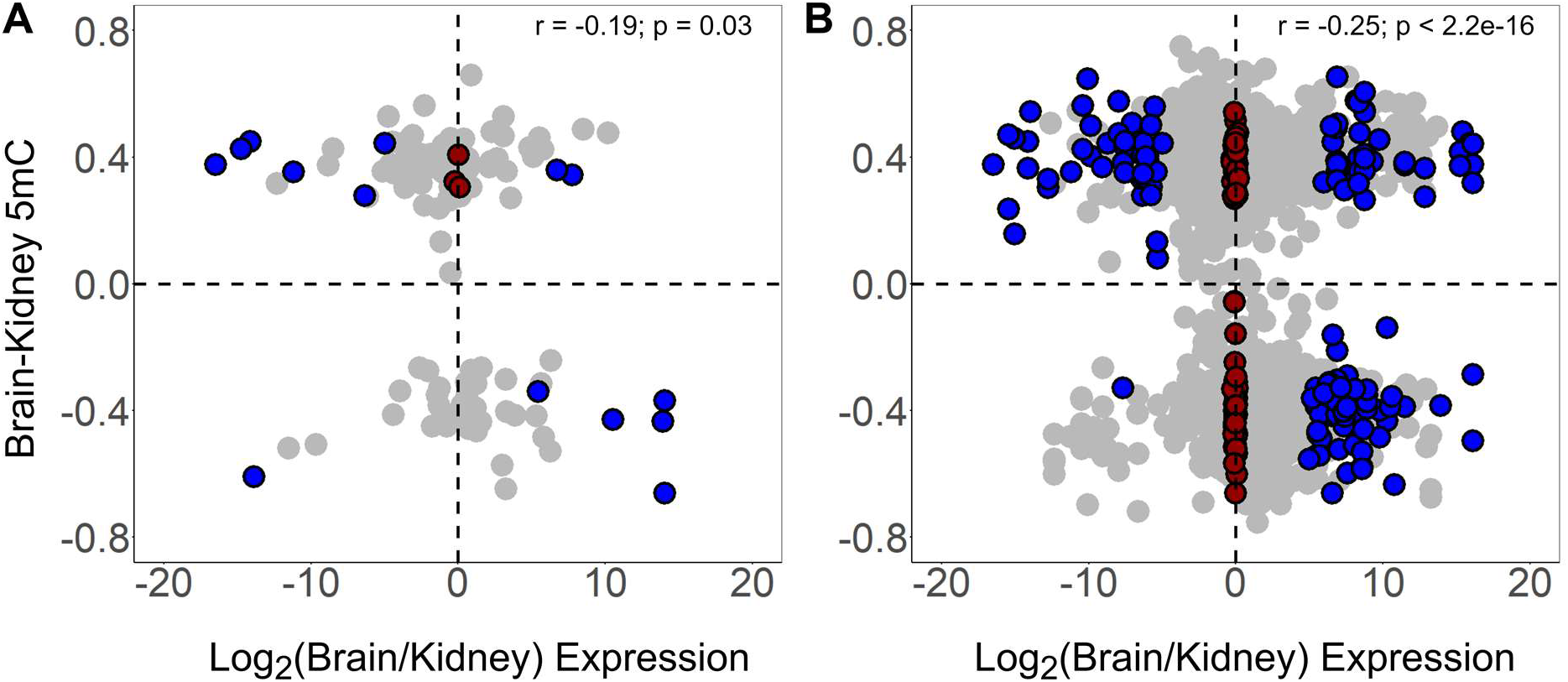
Correlation of tissue dependent DNA methylation (5mC) and gene expression from brain and kidney samples. (A) The mean brain and kidney DNA methylation difference calculated for all CpGs across each gene promoter matched with the corresponding log-transformed ratio of brain to kidney expression. (B) The mean brain and kidney DNA methylation difference calculated for all CpGs across each gene body and matched with corresponding log-transformed ratio of brain to kidney expression. For A and B, Spearman’s rank correlation coefficient and the associated p-value is reported. Blue dots indicate genes that are significantly differentially express between brain and kidney samples (probability of differential expression > 95% based on NOISeq) and red dots show all genes that are significantly similarly methylated in brain and kidney samples (probability of differential expression < 5% based on NOISeq).

**Supplementary Figure 4.**
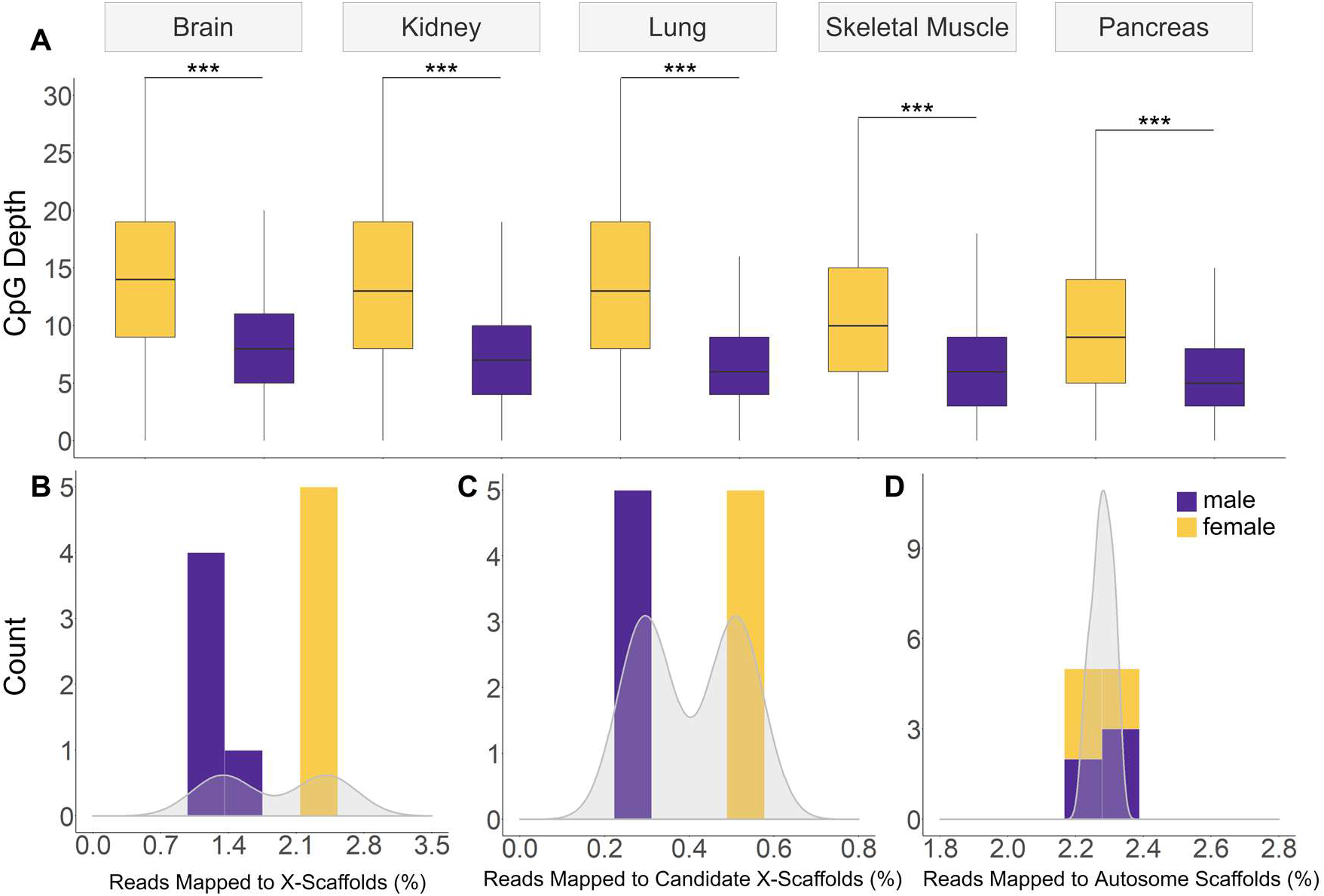
Sex-specific CpG depth of coverage and read mapping to autosomes and X chromosomes. (A) For each of the five tissues, box-and-whisker plots of CpG depths across the X chromosome in male (purple) and female (orange) samples (*** indicates p < 2.2×10-16, Mann-Whitney U test). Histogram and distribution of sex-based read mapping per sample (n=10) to (B) X-linked scaffolds, (C) candidate X-linked scaffolds, and (D) a subset of autosome-linked scaffolds matched in length with all known X-linked scaffolds. For A-C, the percent of reads mapping to the scaffold category of interested over the total number of mapped reads in the genome was calculated for all male (n=5) and female samples (n=5). The known X-linked and candidate X-linked scaffolds show a bimodal distribution with an increase of read mapping to female samples expected from the 2:1 ratio X chromosomes in females to males. This bimodality is not observed in autosomes.

**Supplementary Figure 5.**
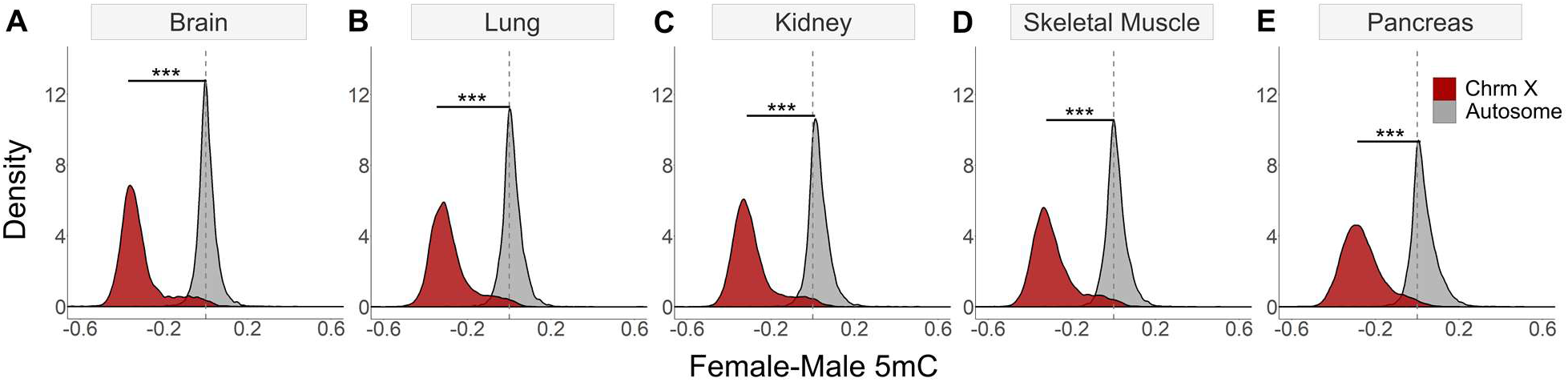
The distribution of sex-based CpG fractional DNA methylation (5mC) differences across autosomes and X chromosomes. The distribution of female and male mean fractional methylation difference from (A) brain, (B) lung, (C) kidney, (D) skeletal muscle and (E) pancreas samples across autosomes and X chromosomes. For A-E, the female and male mean fractional methylation (methylated reads/total reads per CpG) was calculated for all CpGs within 10 kb bins across each autosome-or X-linked scaffold. All tissues exhibited a significant shift towards female hypomethylation in the X chromosome compared to the autosome (*** indicates p < 2.2×10-16, Welch’s t-test).

**Supplementary Figure 6.**
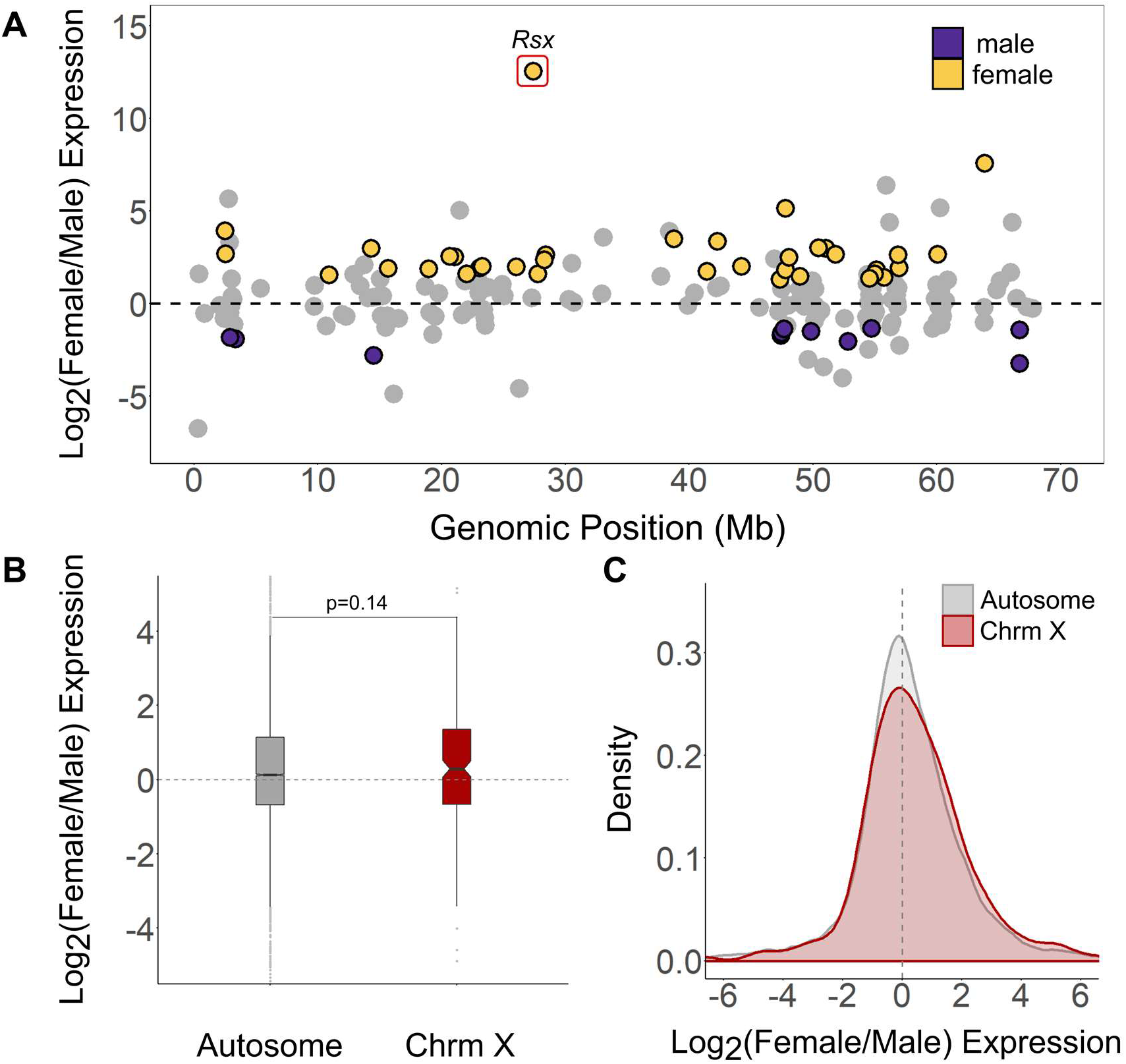
Female and male gene expression across autosomes and the X chromosome using kidney RNA-seq data. (A) The relative genomic location of all genes linked to X chromosome scaffolds aligned by scaffold length and the log-transformed female to male expression ratio generated by NOISeq. Orange dots indicate the 36 genes with significant female-biased expression and purple dots indicate the 11 genes with significant male-biased expression (probability of differential expression > 95% based on NOISeq). For all autosome-linked genes (n = 10,414) and chromosome X-linked genes (n= 209), a box-and-whisker plot (B) and density distribution (C) of the log-transformed female to male expression ratio (p = 0.14, Mann-Whitney U test).

**Supplementary Figure 7.**
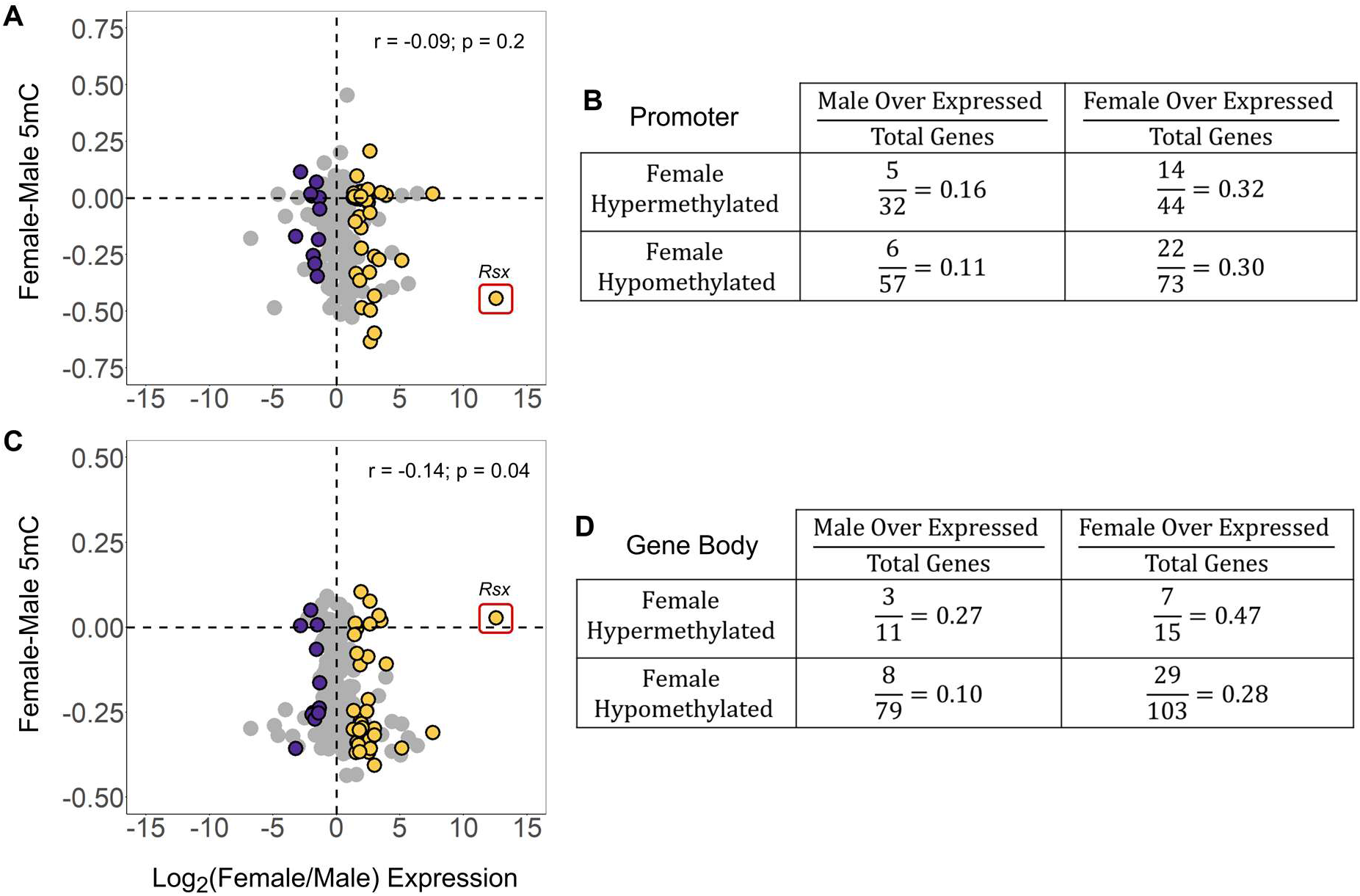
Correlation analysis of sex-based DNA methylation (5mC) and gene expression across chromosome X using kidney WGBS and RNA-seq data. (A) The mean female and male fractional methylation difference calculated for all CpGs across each gene promoter matched with the corresponding log-transformed ratio of female to male expression. (B) The ratio of the number female hypermethylated and female hypomethylated gene promoters that show either significant male of female biased expression over the total number of genes in each category. (C) The mean female and male fractional methylation difference calculated for all CpGs across each gene body and matched with corresponding log-transformed ratio of female to male expression. (D) The ratio of the number female hypermethylated and female hypomethylated gene bodies that show either significant male of female biased expression over the total number of genes in each category. For A and C, Spearman’s rank correlation coefficient and the associated p-value is reported. The Rsx gene was excluded from the correlation calculation. Orange dots indicate the 36 genes with significant female-biased expression and purple dots indicate the 11 genes with significant male-biased expression (probability of differential expression > 95% based on NOISeq).

**Supplementary Figure 8.**
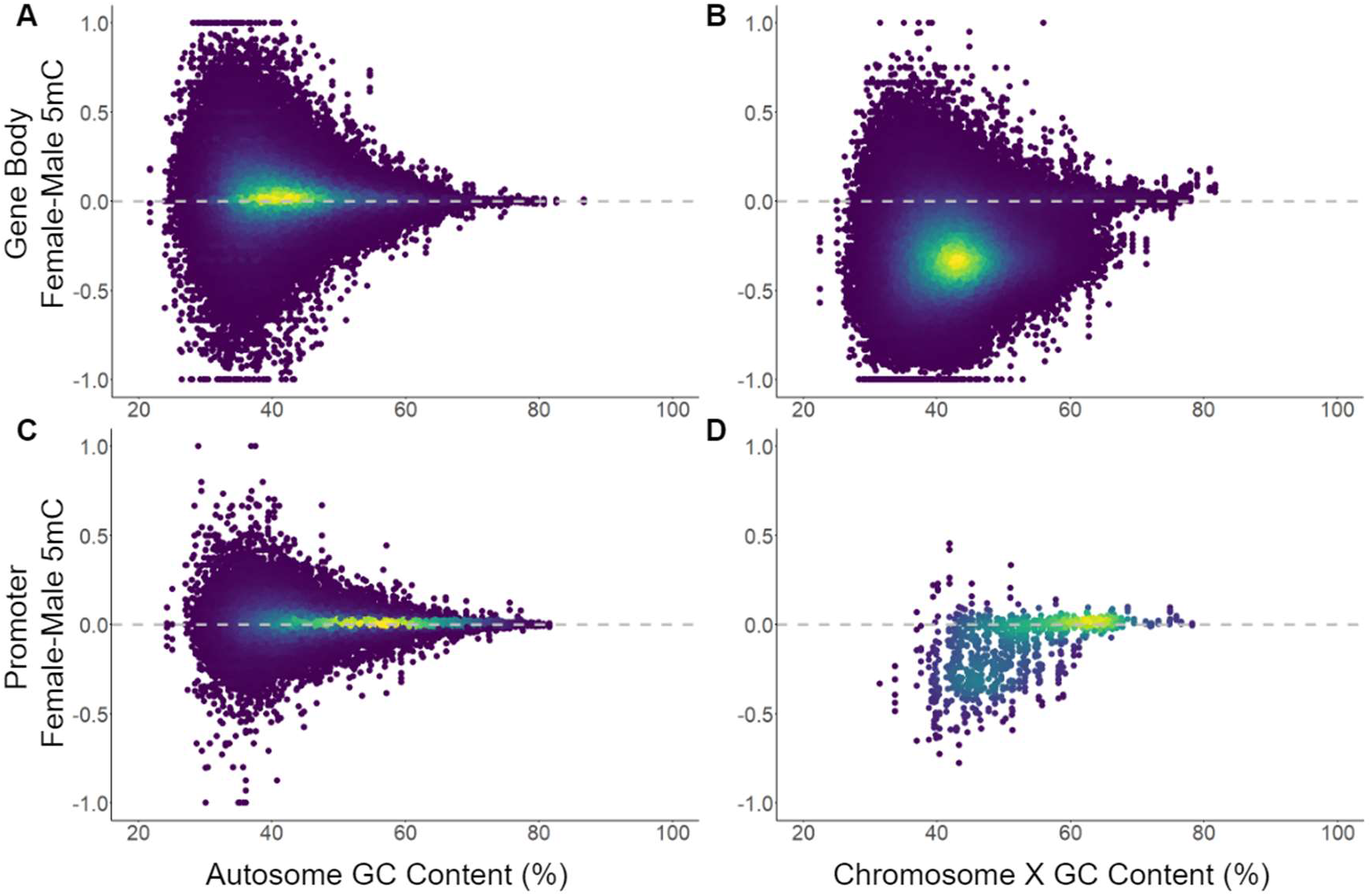
Sex-based DNA methylation (5mC) by GC-content across autosomes and the X chromosome. For A and B, the mean female and male methylation difference calculated from CpGs in 1 Kb bins across (A) autosomes and (B) X chromosomes. For C and D, mean female and male methylation difference calculated from CpGs located in promoter regions (defined as regions 1 kb upstream of known gene TSSs) in (C) autosomes and (B) X chromosomes. For A-D, data from all five tissues (brain, kidney, lung, pancreas, and skeletal muscle) are reported. All plots are coloured by data density where blue represents low density regions and yellow represents high density regions.

